# Exploring recurrent urinary tract infection: paired microbiota and culture-based profiling reveals stable dysbiosis and multidrug-resistant, virulence-enriched uropathogens

**DOI:** 10.64898/2026.10.10.758225

**Authors:** Saumya Kaushik, P Karthika, Surajit Pal, Nancy Nandi, George Seghal Kiran, S. Hari Krishna Kumar, Joseph Selvin, Vajid Nettoor Veettil, Sheela Devi

## Abstract

**Objectives:** Recurrent urinary tract infection (rUTI) is increasingly complicated by antimicrobial resistance (AMR), yet urinary microbiome dysbiosis and isolate-level resistance-virulence are rarely examined together across different treatment states. In this study we characterized both within the same sample in rUTI and in healthy controls.

**Methods:** Cross-sectional cohort, mid-stream urine was collected from women with untreated, treated and prophylaxis rUTI and age-matched healthy controls at four centres in South India. Of 80 samples collected, 75 were analysed (26 controls, 20 untreated, 20 treated, 9 prophylaxis) by 16S rRNA V3-V4 sequencing; in addition, 105 culturable isolates from the same samples were identified and tested for antimicrobial susceptibility (VITEK 2) and phenotypic virulence.

**Results:** rUTI groups showed approximately half the alpha diversity of controls (median Shannon 1.07-1.18 vs 2.17; Kruskal-Wallis *p* = 0.041). Community structure differed from controls in every rUTI sub-group (PERMANOVA R^2^ = 0.091, *p* = 0.004; pairwise FDR = 0.008) but not among sub-groups. Mean *Lactobacillus* relative abundance fell from 29.2% in controls to ≤13.2% in rUTI groups, while *Escherichia*-*Shigella* rose from 7.2% to 25.0-43.0%. Control-referenced dysbiosis z-scores were elevated in all rUTI groups (community distance: FDR ≤ 0.003; uropathogen: *Lactobacillus* index: FDR < 0.001 for treated and prophylaxis groups). Among isolates, *Escherichia coli* predominated (35/105; 16/35, 45.7% multidrug-resistant), followed by *Klebsiella* spp. (32% multidrug-resistant); carbapenem susceptibility was largely retained, and multidrug-resistant *E. coli* co-carried more virulence traits.

**Conclusions:** rUTI is associated with a stable dysbiotic state that treatment does not restore, combining *Lactobacillus* loss and uropathogen dominance with a multidrug-resistant culturable reservoir that co-accumulates virulence traits. Reading the community and its isolates together may support microbiome-sparing, resistance-aware management of rUTI.

## Introduction

Urinary tract infection (UTI) is among the most prevalent microbial infections worldwide, reporting 400 million incident cases estimated annually [1]. A substantial proportion of affected women experience recurrence: recurrent UTI (rUTI), defined as two or more culture-proven incidences occurring within six months or three or more incidences within twelve months, requires repeated antibiotic courses and remains difficult to manage [2,3].

The clinical course of rUTI spans distinct states. Untreated rUTI is sustained by quiescent intracellular bacterial reservoirs that repeatedly seed the bladder lumen [4]; treated rUTI reflects persistence or reinfection after completed therapy; and prophylaxis rUTI occurs despite long-term low-dose suppressive antibiotics [5]. Each state imposes differential antibiotic selective pressure that favors multidrug-resistant (MDR) uropathogens, and rising resistance to β-lactams, cephalosporins, fluoroquinolones and carbapenems increasingly constrains management [6,7]. In parallel, uropathogen fitness depends on virulence determinants adhesive fimbriae, motility, biofilm formation, nutrient acquisition and extracellular enzymes whose relationship with resistance is incompletely understood [7].

Culture-independent profiling has shown that the healthy female urinary microbiota is frequently *Lactobacillus*-rich and that dysbiosis commensal loss with uropathogen expansion which accompanies rUTI [8,9]. Sequencing alone, however, cannot say which organisms are viable, which carry resistance determinants, or which express the virulence machinery that drives tissue invasion; culture, conversely, recovers only the fraction of the community that grows on a plate, stripped of ecological context. Reading the same specimens through both lenses is more than the sum of the parts: the community map shows where each isolate sits ecologically, while isolate phenotypes attach clinical meaning such as treatability and pathogenic potential to taxa that sequencing can only name. Few rUTI studies have pursued both levels in depth, and fewer still have asked how the resistance burden of the culturable reservoir relates to the virulence it carries, although that relationship decides whether clearing dysbiosis also defuses it. How dysbiosis, and the resistance-virulence balance of the isolates it harbours, differ across untreated, treated and prophylaxis states remains poorly defined.

We therefore performed a cross-sectional cohort study combining 16S rRNA V3-V4 sequencing of 75 mid-stream urine samples with antimicrobial susceptibility and phenotypic virulence testing of 105 culturable isolates from the same specimens. Our objectives were to (i) compare diversity, composition and control-referenced dysbiosis scores across rUTI treatment states and healthy controls, and (ii) define the resistance and virulence profiles of the culturable uropathogen reservoir, including resistance-virulence co-occurrence.

## Methods

Study design and participants: The study is reported following the Strengthening the Reporting of Observational Studies in Epidemiology (STROBE) guideline. Women with rUTI (untreated, treated, or receiving antibiotic prophylaxis) and age-matched healthy controls were enrolled at IQRAA Hospital (Kerala), Auroville Health Centre (Tamil Nadu), Pondicherry Institute of Medical Sciences and the local Pondicherry community during 2024-25. Inclusion required ≥2 culture-proven symptomatic UTIs within 6 months or ≥3 within 12 months. Exclusion criteria were: unwillingness to discontinue UTI-prevention supplements (methenamine, probiotics, D-mannose, cranberry) 6 weeks before sampling; urinary tract instrumentation within the previous 3 days; neurogenic bladder or chronic intermittent catheterisation; prior pelvic radiation; active malignancy, including breast cancer within the previous five years; neurologic conditions affecting urinary function (stroke, multiple sclerosis, spinal cord injury, Parkinson’s disease); renal insufficiency (creatinine >1.3 mg/dL); immunosuppression or chronic steroid use; intravaginal pessary use; post-coital antibiotic prophylaxis (excluded from all sub-groups); antibiotics for any active infection within the previous month; and anti-oestrogen medication. Ethics approval was obtained from the Institutional Ethical Committee (Human Studies), Pondicherry University; all participants gave written informed consent, and demographic and medical histories were recorded.

Sample collection and processing: Mid-stream clean-catch urine (20-30 mL) was collected aseptically after cleansing of the external genitalia and discarding the initial stream, transported chilled, and processed promptly: one aliquot for culture and remaining for DNA extraction. DNA was extracted using DNeasy Blood & Tissue Kit (Qiagen) with proteinase K lysis, quantified by Qubit fluorometry, assessed by A260/A280 ratio and agarose gel electrophoresis, and stored at -80°C.

16S rRNA sequencing and bioinformatics: Amplification and one paired end sequencing was carried out for 16S rRNA V3-V4 region. Reads were quality-filtered (Phred ≥30; FastQC) and adapter-trimmed (fastp), then processed in QIIME 2 with OTU clustering at 97% identity against SILVA v138.2 [10-12]. Analyses used R 4.4.2 with vegan 2.7.6 and phyloseq [13,14]. Alpha diversity (Shannon, Simpson, inverse Simpson, observed richness, Pielou evenness) was compared by Kruskal-Wallis tests with pairwise Wilcoxon follow-up (Benjamini-Hochberg FDR). Beta diversity used Bray-Curtis dissimilarities: NMDS (metaMDS), PERMANOVA (adonis2, 999 permutations) with betadisper dispersion checks, and pairwise PERMANOVA (FDR). Two control-referenced dysbiosis scores were computed per sample and z-standardised to the control distribution (mean 0, SD 1). Score A is the Bray-Curtis distance to the healthy-control centroid i.e, how far the community has drifted from the healthy reference. Score B, the uropathogen:Lactobacillus index, is log10((U + 10^−4^)/(L + 10^−4^)), where U is the summed relative abundance of twelve uropathogenic genera (*Escherichia*-*Shigella, Klebsiella, Proteus, Enterococcus, Pseudomonas, Staphylococcus, Streptococcus, Serratia, Acinetobacter, Morganella, Providencia, Citrobacter*) and L is *Lactobacillus* relative abundance. The index reads intuitively: 0 indicates equal uropathogen and *Lactobacillus* abundance, +1 a ten-fold uropathogen excess and −1 a ten-fold *Lactobacillus* excess; the 10^−4^ pseudocount keeps the ratio defined when either partner is undetected. Because both scores are z-standardised, a value of 0 equals the healthy mean, 1 lies one standard deviation above it, and |z| > 1.96 falls outside the healthy 95% reference range. Urotypes were assigned by each sample’s dominant genus and compared by Fisher’s exact test. Genus-level differential abundance (pooled rUTI versus controls) used Wilcoxon tests with FDR correction.

Isolate characterization: Culturing of samples were done on cysteine-lactose-electrolyte-deficient agar (37°C, 24 h) [15]; morphologically distinct colonies were purified, glycerol-stocked, and identified with antimicrobial susceptibility testing on the VITEK 2 Compact system (bioMérieux, Marcyl’Étoile, France) as per manufacturer’s instructions. MDR was defined as resistance to ≥4 tested agents. Phenotypic virulence assays comprised urease production, lipase production (rhodamine), starch hydrolysis, citrate utilisation, haemolysis on sheep blood agar, biofilm formation on sucrose-supplemented Congo red brain-heart infusion agar, and swimming/swarming motility.

## Results

Cohort: Eighty samples were collected; five (three controls and 2 untreated rUTI) failed DNA quality control, leaving 75 analysed: 26 controls, 20 untreated, 20 treated and 9 prophylaxis rUTI (participant flow and demographics, Supplementary Table S1).

Urinary microbiome diversity and composition: Alpha diversity was highest in controls (median Shannon 2.17, IQR 1.39; median observed richness 114 genera, IQR 79) and roughly halved across rUTI groups (Shannon 1.07-1.18; richness 56-75.5; Kruskal-Wallis *p* = 0.041 and p = 0.024, respectively; Fig. 1A,B; Table 1). Simpson and inverse-Simpson indices showed the same pattern (median Simpson 0.73 in controls vs 0.39-0.42 in rUTI; p = 0.037), whereas Pielou evenness did not differ significantly (*p* = 0.075). Pairwise control-versus-rUTI Wilcoxon tests were nominally significant (e.g., control vs untreated richness *p* = 0.006) but did not survive FDR correction. Bray-Curtis NMDS separated controls from rUTI samples (stress = 0.197; Fig. 1C); global PERMANOVA confirmed the group effect (R^2^ = 0.091, *p* = 0.004; betadisper *p* = 0.236, indicating homogeneous dispersions), and every rUTI sub-group differed from controls (pairwise PERMANOVA FDR = 0.008) while sub-groups did not differ from each other (FDR > 0.14). Compositionally, controls were *Lactobacillus*-rich (mean 29.2%) with low *Escherichia*-*Shigella* (7.2%), whereas rUTI groups were dominated by *Escherichia*-*Shigella* (untreated 25.0%, treated 37.1%, prophylaxis 43.0%) with *Klebsiella* enrichment (9.6-11.6% vs 2-3% in controls) and *Lactobacillus* depletion (13.2%, 7.7% and 1.1%, respectively; Fig. 1D). Phylum-level Proteobacteria expanded from 45.0% (controls) to 61.8-77.3% (rUTI) at the expense of Firmicutes, and Bacteroidota and Fusobacteriota contracted in parallel (Supplementary Fig. S1). *Gardnerella* remained detectable across groups (mean 3.8-4.2%), a potential uropathogen reservoir implicated in reactivation of dormant *Escherichia coli* [16]. Per-sample profiles confirmed group-level shifts with substantial inter-individual heterogeneity (Supplementary Fig. S2).

**Table 1.** Diversity, composition and dysbiosis metrics by study group.

| Group | n | Shannon,<br>median<br>(IQR) | Observed<br>richness,<br>median<br>(IQR) | Lactobacillus,<br>mean % | Escherichia-Shigella,<br>mean % | Distance<br>z,<br>median | PERMANOVA<br>vs control,<br>FDR |
| --- | --- | --- | --- | --- | --- | --- | --- |
| Healthy<br>control | 26 | 2.17<br>(1.39) | 114 (79) | 29.2 | 7.2 | 0.09 | - |
| Untreated<br>rUTI | 20 | 1.16<br>(1.42) | 56 (41) | 13.2 | 25.0 | 1.52 | 0.008 |
| Treated<br>rUTI | 20 | 1.07<br>(1.71) | 75.5<br>(68.5) | 7.7 | 37.1 | 1.43 | 0.008 |
| Prophylaxis<br>rUTI | 9 | 1.18<br>(1.63) | 64 (29) | 1.1 | 43.0 | 1.96 | 0.008 |

IQR, interquartile range; rUTI, recurrent urinary tract infection. Kruskal-Wallis across groups: Shannon $p = 0.041$ ; richness $p = 0.024$ . PERMANOVA on Bray-Curtis dissimilarities, Benjamini-Hochberg-adjusted.

**Fig. 1.**
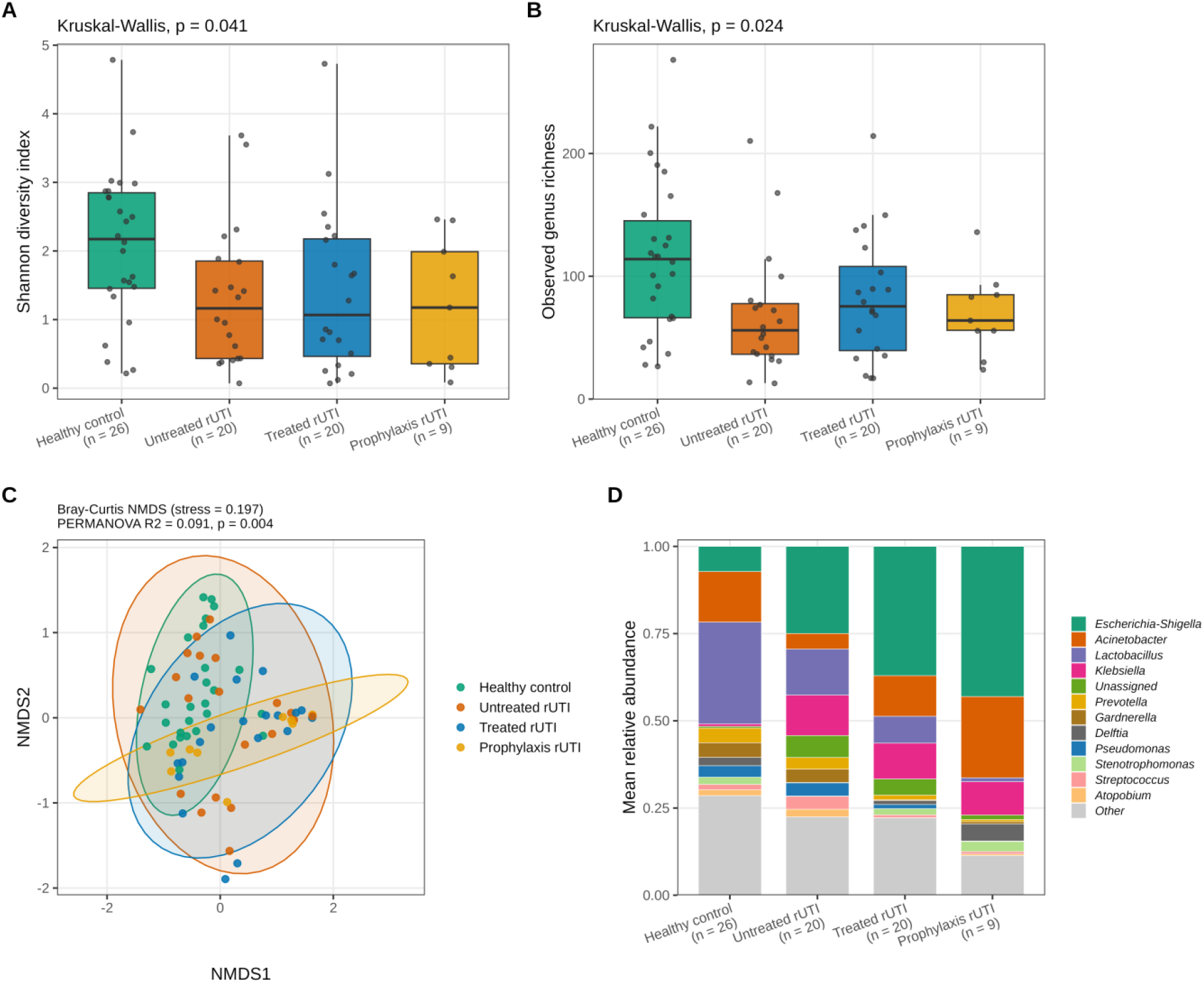
Urinary microbiome diversity and composition across rUTI sub-groups and healthy controls. (A) Shannon diversity and (B) observed genus richness per group (boxplots with per-sample points; Kruskal-Wallis p shown; no pairwise Wilcoxon comparison survived Benjamini-Hochberg correction). (C) Bray-Curtis NMDS ordination (stress = 0.197) with 95% confidence ellipses; global PERMANOVA R^2^ = 0.091, *p* = 0.004; all control-versus-rUTI pairwise PERMANOVAs FDR = 0.008. (D) Mean genus-level relative abundance per group (top 12 genera + Other). C, healthy control (n = 26); rN, untreated rUTI (n = 20); rT, treated rUTI (n = 20); rP, prophylaxis rUTI (n = 9).

Control-referenced dysbiosis scores and urotypes: Both dysbiosis scores were elevated in rUTI (Fig. 2). Community distance to the healthy reference was higher in all rUTI groups (median z 1.43-1.96 vs 0.09; FDR ≤ 0.003), and the uropathogen: *Lactobacillus* index was elevated in treated and prophylaxis groups (median z 1.13-1.20; FDR < 0.001) but not untreated (FDR = 0.082); in practical terms, the typical treated or prophylaxis sample carried a uropathogen-versus-*Lactobacillus* balance more than one standard deviation above the healthy mean. Urotype distribution differed between controls and pooled rUTI (Fisher exact p < 0.001): 11/26 controls (42%) were *Lactobacillus*-dominant versus 4/49 rUTI samples (8%), while *Escherichia*-*Shigella*-dominant urotypes occurred in 2/26 controls (8%) versus 17/49 rUTI samples (35%) (Supplementary Fig. S3).

**Fig. 2.**
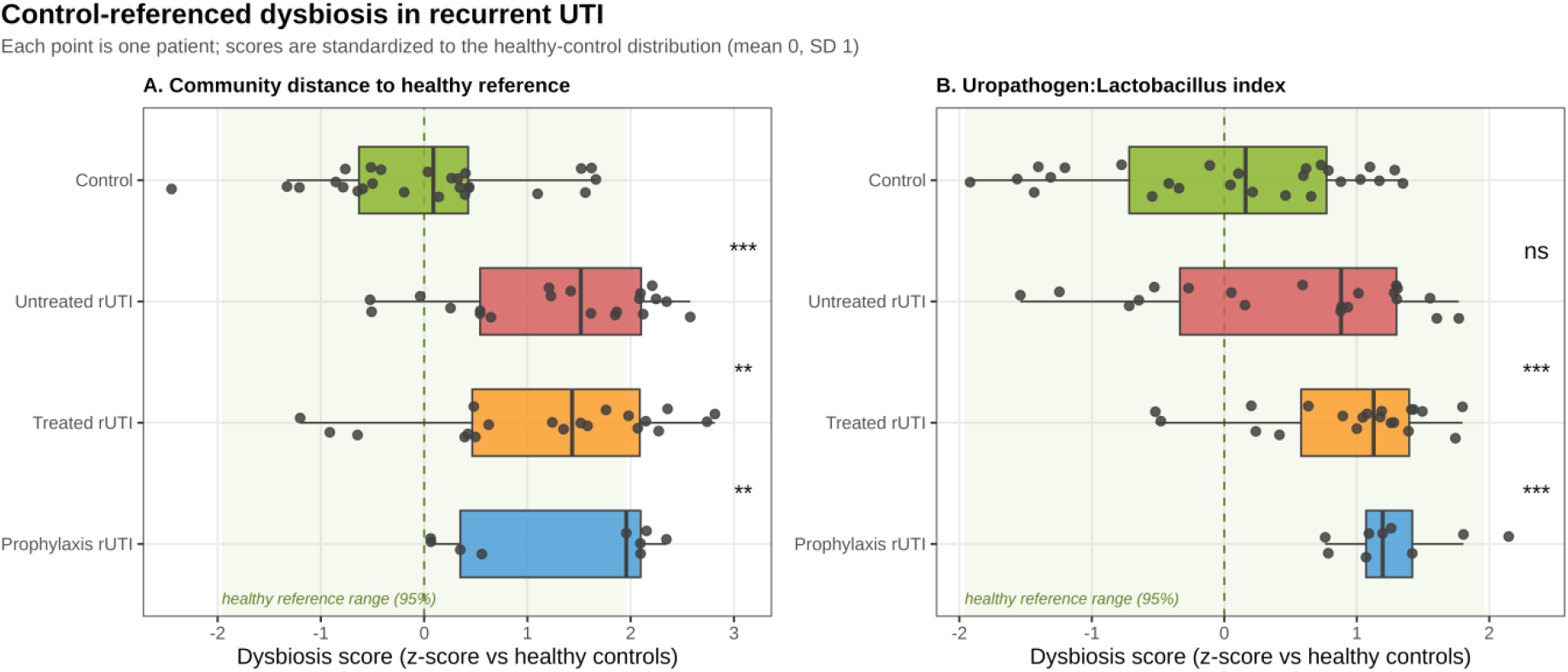
Control-referenced dysbiosis scores. Each point is one participant; scores are z-standardised to the healthy-control distribution (mean 0, SD 1); shaded band, healthy 95% reference range. (A) Bray-Curtis distance to the healthy-control centroid. (B) Uropathogen:Lactobacillus index (log10 ratio). Wilcoxon tests versus controls, Benjamini-Hochberg-adjusted: **FDR < 0.01, ***FDR < 0.001; ns, not significant.

Resistance and virulence of culturable uropathogens: Urine culture yielded 105 uropathogenic isolates; 76 were confidently identified and 66 had complete susceptibility profiles; the remainder were unidentifiable or low-discrimination by VITEK 2 (Supplementary Table S2). *Escherichia coli* predominated (35 isolates; 16/35, 45.7% MDR), followed by the *Klebsiella* complex (*K. pneumoniae, K. pneumoniae* subsp. *pneumoniae* and *ozaenae, K. aerogenes*; 32% MDR); *Serratia ficaria, Serratia fonticola, Proteus mirabilis* and *Sphingomonas paucimobilis* were isolated less frequently (Fig. 3). Resistance to penicillins was widespread in *E. coli* and *Klebsiella*, whereas the β-lactamase-inhibitor combinations piperacillin-tazobactam and cefoperazone-sulbactam remained active against most isolates, with resistance restricted to a few MDR strains. Cephalosporin activity improved with generation, first-generation agents faced broad resistance while later generations retained higher efficacy and carbapenem susceptibility was largely preserved, with resistance confined to a few *Klebsiella* isolates. Most isolates hydrolysed starch, utilised citrate, formed biofilm and were motile, consistent with fimbriae expression and biofilm-mediated colonisation; advanced traits (extracellular lipase, α-haemolysis) clustered in MDR *E. coli*, whereas *Klebsiella* isolates compensated for absent motility with citrate utilisation and lipase activity (Fig. 4). The resistance-virulence relationship was not uniform across taxa: positive in *E. coli*, static in *Klebsiella* and inverse in *Serratia*.

**Fig. 3.**
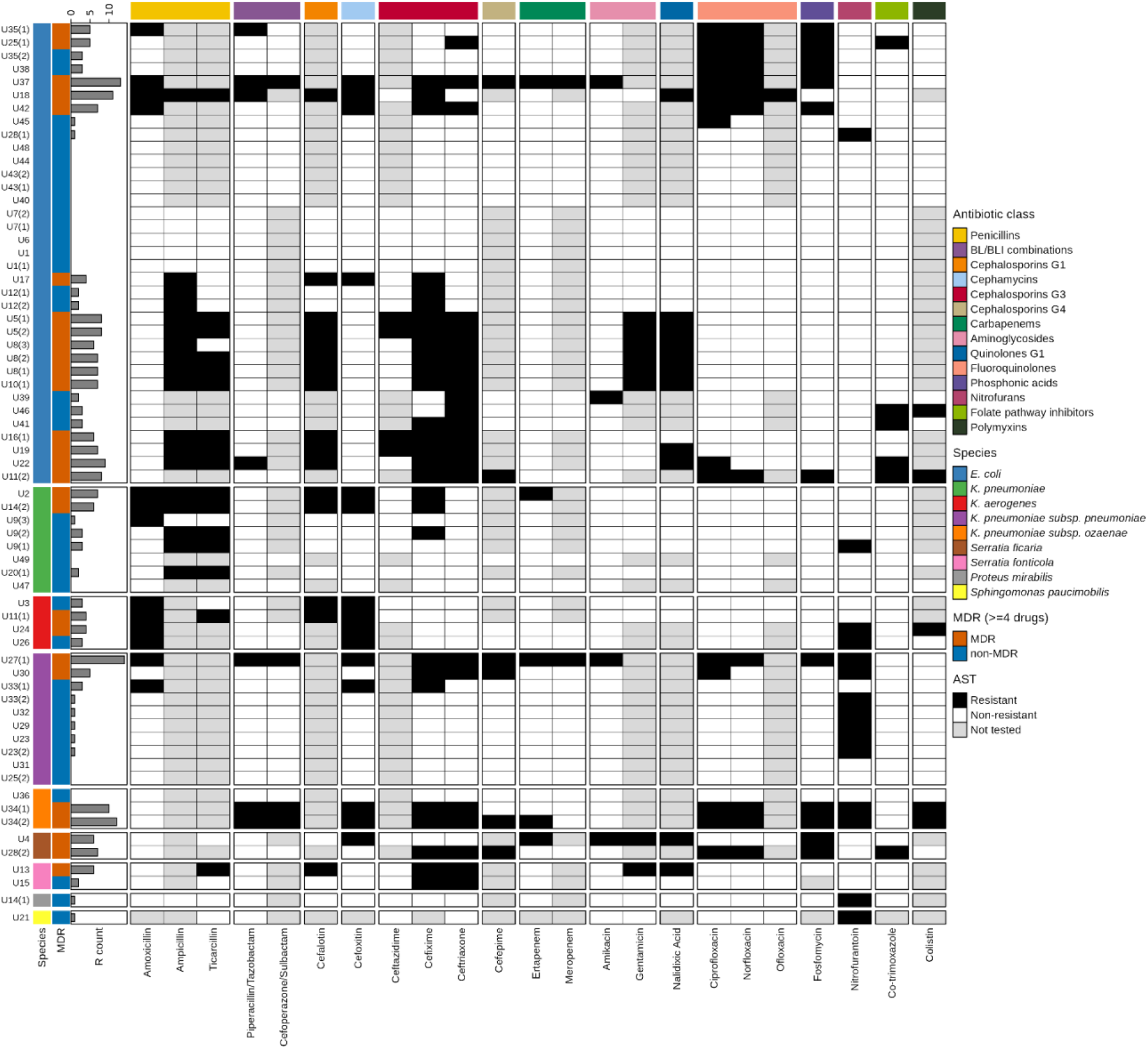
Antimicrobial susceptibility profiles of culturable urinary isolates. Heatmap of isolate-level susceptibility (black, resistant) across antibiotic classes, with species and multidrug-resistance (≥4 agents) annotation; inset, MDR versus non-MDR counts by species.

**Fig. 4.**
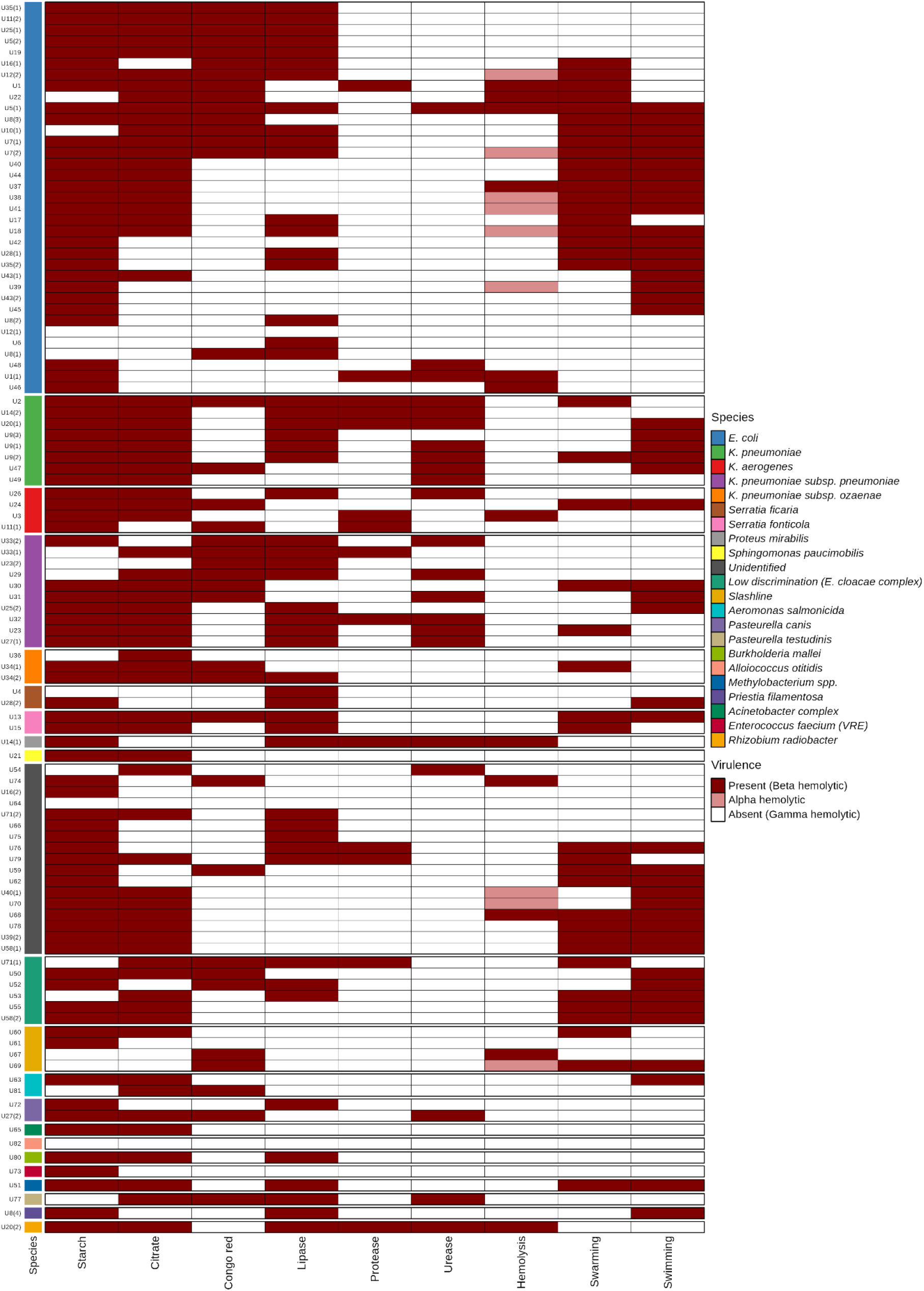
Phenotypic virulence profiles of culturable urinary isolates. Heatmap of nine virulence assays (starch hydrolysis, citrate utilisation, biofilm, lipase, protease, urease, haemolysis, swarming and swimming motility) across isolates, annotated by species and MDR status; advanced traits (lipase, α-haemolysis) cluster in MDR *Escherichia coli*.

## Discussion

Principal findings: By pairing 16S sequencing with culture-based profiling of the same mid-stream specimens, this study examined recurrent UTI at two levels at once. The community view showed a stable dysbiotic state: alpha diversity roughly halved, composition separated from healthy controls in all three treatment states, *Lactobacillus* depleted in step with antibiotic pressure, and uropathogenic genera dominant. The isolate view showed what that dominance means in practice a culturable reservoir led by multidrug-resistant *E. coli* and *Klebsiella*, with advanced virulence traits concentrated in the most resistant *E. coli*. Neither readout alone would have exposed how tightly the ecological shift and the clinical threat are coupled.

Comparison with previous work: Reduced urinary diversity and compositional separation from controls agree with previous 16S studies of rUTI cohorts [17-19]. Striking here is the gradient: mean *Lactobacillus* abundance fell stepwise from 29.2% in controls to 13.2% (untreated), 7.7% (treated) and 1.1% (prophylaxis), tracking cumulative antibiotic exposure more closely than clinical category. The near-eradication of *Lactobacillus* under prophylaxis mirrors reports that daily suppressive antibiotics deplete protective lactobacilli [20] and that oestrogen-associated *Lactobacillus* enrichment tracks with reduced rUTI susceptibility [19]; mechanistically, *L. crispatus* restricts bladder uropathogenic *E. coli* through a host type-I interferon response [9], so its loss removes both microbial and immunological restraint. Equally telling is what treatment did not do: treated and prophylaxis communities remained control-distinct and indistinguishable from untreated rUTI, suggesting that therapy resolves symptoms without rebuilding the community consistent with prophylaxis trials in which protection lasts only while the drug is taken [5,21] and with evidence that sub-inhibitory exposure can itself remodel uropathogen virulence and host immunity [7].

The isolate data speak to a dilemma at the heart of rUTI management: Does selecting for resistance also select for virulence? Our MDR frequencies (45.7% of *E. coli*) align with Indian and international uropathogen surveys [22-24], and the positive resistance-virulence coupling in *E. coli* supports co-selection, potentially through plasmid co-carriage of resistance and virulence loci [25,26]. But the coupling was not universal rather it was static in *Klebsiella* and inverse in *Serratia* and experimental evolution tells a similarly drug-specific story: in *Pseudomonas aeruginosa* evolved under single antibiotics, ciprofloxacin resistance increased virulence, piperacillin-tazobactam resistance reduced virulence through genomic deletion, and streptomycin resistance was roughly neutral [27]. Read together, these patterns argue against any universal resistance-virulence trade-off; the direction depends on the organism and, critically, on the drug. If that holds in the urinary tract, antibiotic choice in rUTI may shape not only tomorrow’s resistance but also the virulence of the reservoir that survives therapy favouring agents whose resistance carries a virulence cost, and urging caution where resistance and virulence rise together, as reported for fluoroquinolones [27]. Our cross-sectional phenotypes cannot test this directly, but they make the question concrete.

Strengths and limitations: Strengths include paired community- and isolate-level profiling of the same specimens, four clinically defined groups spanning the treatment course, multi-centre enrolment, and control-referenced dysbiosis scores that place each patient on a healthy-referenced scale. Limitations temper interpretation. The cross-sectional design precludes causal inference: dysbiosis may predispose to rUTI, result from it, or both. The prophylaxis arm was small (n = 9). Relative-abundance data preclude absolute quantification and rarefaction-based richness estimators, and genus-level V3-V4 resolution cannot distinguish uropathogenic from commensal *E. coli* strains. Virulence was assayed as in vitro phenotypes that may not reflect expression in-vivo, and our MDR definition (resistance to ≥4 tested agents) is stricter than standard class-based definitions, so frequencies are not directly comparable across studies. Finally, VITEK 2 left 29 of 105 isolates unidentified, and reagent contamination cannot be fully excluded in low-biomass urine sequencing.

Mechanisms: The dysbiosis signature was community-level: no single genus survived FDR correction, yet multivariate and urotype analyses were strongly significant. rUTI thus resembles an alternative stable state of uropathogen bloom with commensal collapse that antibiotics alone do not reset [8,28]. *Gardnerella*, detectable across groups, can reactivate dormant intracellular *E. coli* reservoirs [4,16], offering one route by which a dysbiotic community perpetuates recurrence. Co-accumulation of resistance and virulence in MDR *E. coli* may help explain its long-standing predominance, whereas the inverse coupling in *Serratia* fits a resistance-fitness trade-off in opportunists [25].

Implications for practice and research: Community-level readouts such as: dysbiosis scores, urotypes, the uropathogen: *Lactobacillus* index could complement culture for risk stratification, flagging patients whose microbiome remains control-distinct after treatment and who may benefit from restorative rather than suppressive strategies [28]. Retained carbapenem and β-lactamase-inhibitor combination activity supports local antibiogram-guided therapy and carbapenem stewardship, in line with international guidance on antimicrobial-resistant Gram-negative infections [24,29]. The drug-specific resistance-virulence relationship adds a further consideration: where a choice between active agents exists, their likely evolutionary side-effects on virulence deserve a place in the decision. Longitudinal cohorts with strain-resolved genomics and absolute quantification are needed to test whether dysbiosis precedes recurrence and to track resistance-virulence co-evolution [30].

## Conclusions

Recurrent UTI in this cohort was a disorder of both community and pathogen: a stable, treatment-resistant dysbiosis inhabited by multidrug-resistant isolates carrying concentrated virulence potential. Managing rUTI as an ecological as well as a microbiological problem and choosing antibiotics with an eye on the virulence they select may improve on symptom-by-symptom suppression.

## Acknowledgements

The authors are grateful for the support and cooperation of IQRAA Center for Research and Development, Calicut, Kerala, India, Pondicherry Institute of Medical Sciences, Puducherry, India and Auroville Health Centre Tamil Nadu, India in clinical sample collection for the purpose of the study.

## Data availability

All the data can be made accessible upon request to corresponding author

## Ethical Approval

Approval No. HECPU/2023/0-ER/I/17-10-2023) the research proposal titled: “Dissecting the role of Vaginal and Urinary Tract Microbiome Dysbiosis in Recurrent Urinary Tract Infections and Emergence of Antimicrobial Resistance: A Case Control Study”

## Author contributions

Saumya Kaushik, Karthika P. and Surajit Pal: Literature survey, writing, editing; Saumya Kaushik, Karthika P., Nancy Nandi: Laboratory experiments and validation; Surajit Pal, S. Hari Krishna Kumar: Performed in silico analysis, George Seghal Kiran, Joseph Selvin: Conceptualization, reviewing, editing and approved for final manuscript; Saumya Kaushik, Karthika P., Vajid Nettoor Veettil, Sheela Devi: Sample collection.

---

**Supplementary Fig. S1.**
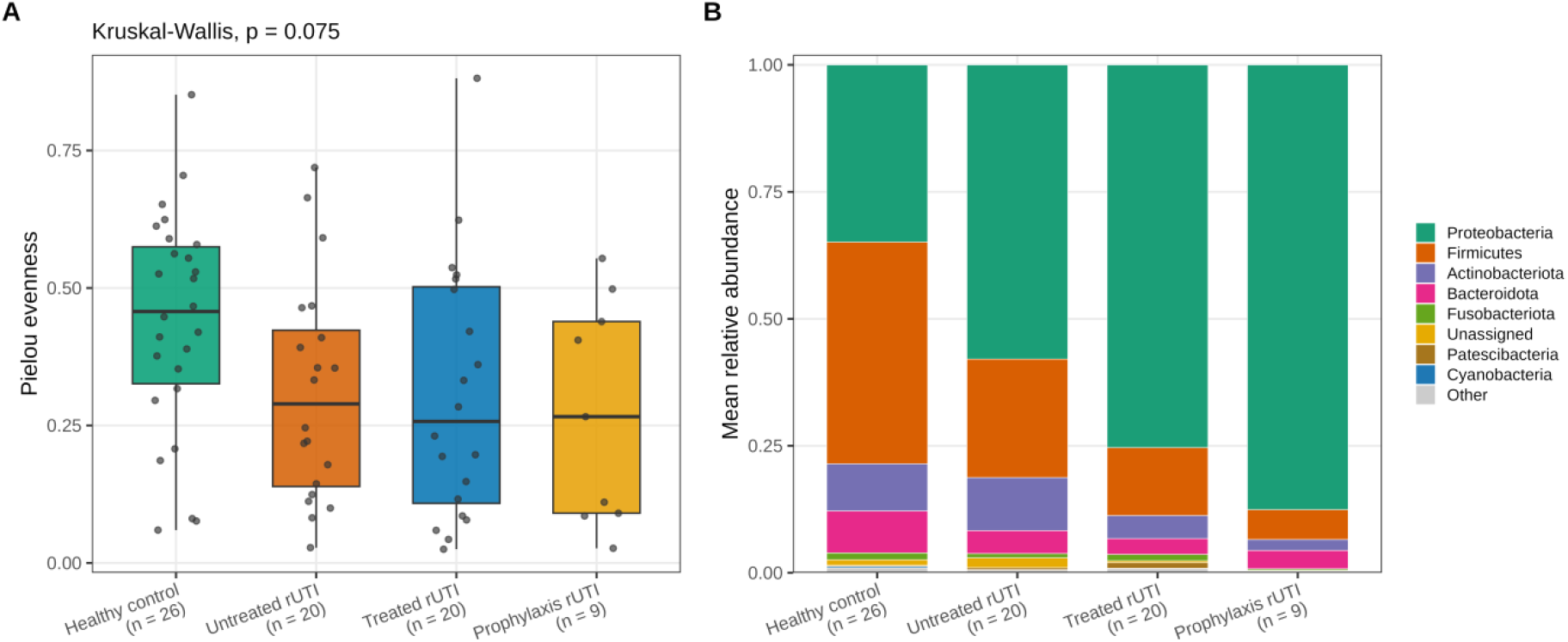
Community evenness and phylum-level composition. (A) Pielou evenness per group (boxplots with per-sample points; Kruskal-Wallis *p* = 0.075). (B) Mean phylum-level relative abundance per group; Proteobacteria expanded from 45.0% in controls to 61.8-77.3% across rUTI groups at the expense of Firmicutes, while Bacteroidota and Fusobacteriota contracted. C, healthy control (n = 26); rN, untreated rUTI (n = 20); rT, treated rUTI (n = 20); rP, prophylaxis rUTI (n = 9).

**Supplementary Fig. S2.**
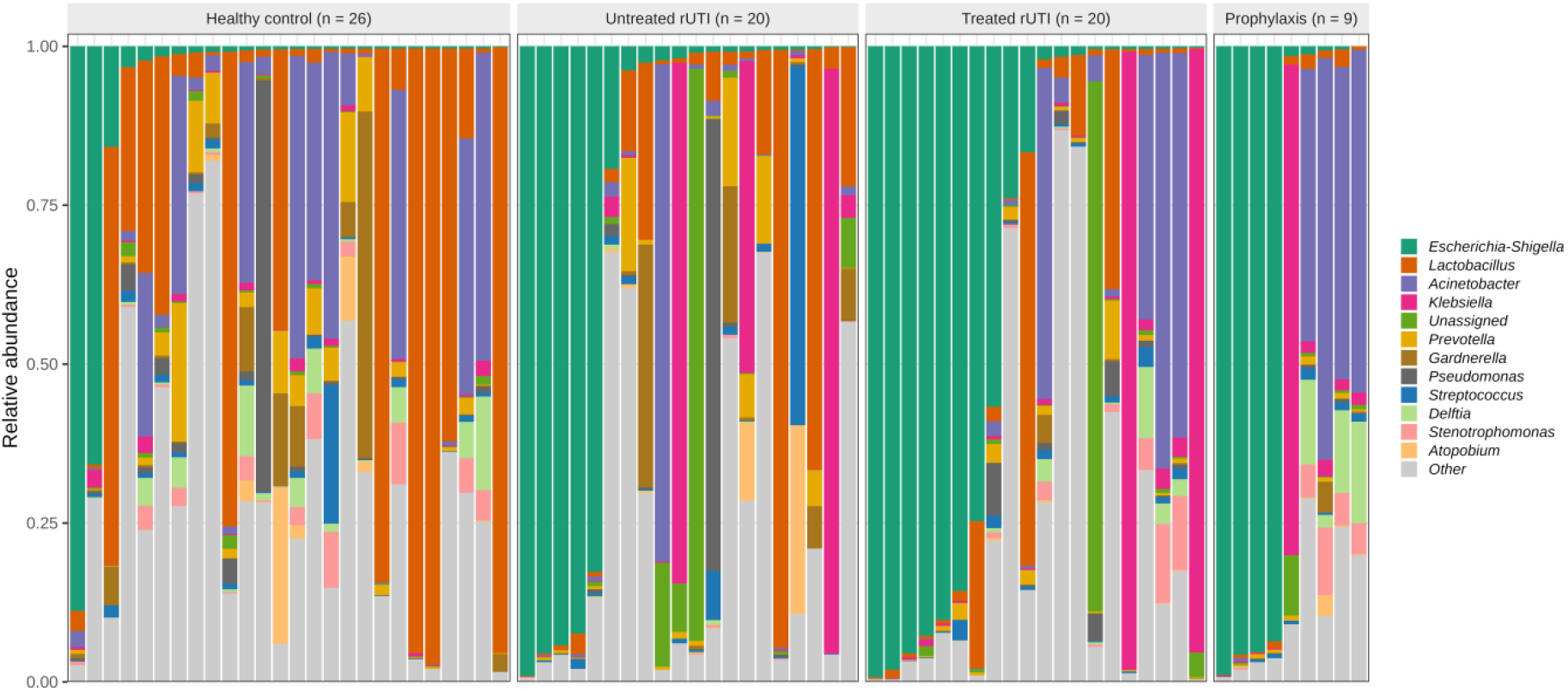
Per-sample genus-level composition of all 75 mid-stream urine samples. Each bar is one sample, grouped by study arm; the most abundant genera are shown individually with the remainder grouped as Other. Group-level shifts towards *Escherichia*-*Shigella* dominance are visible alongside substantial inter-individual heterogeneity.

**Supplementary Fig. S3.**
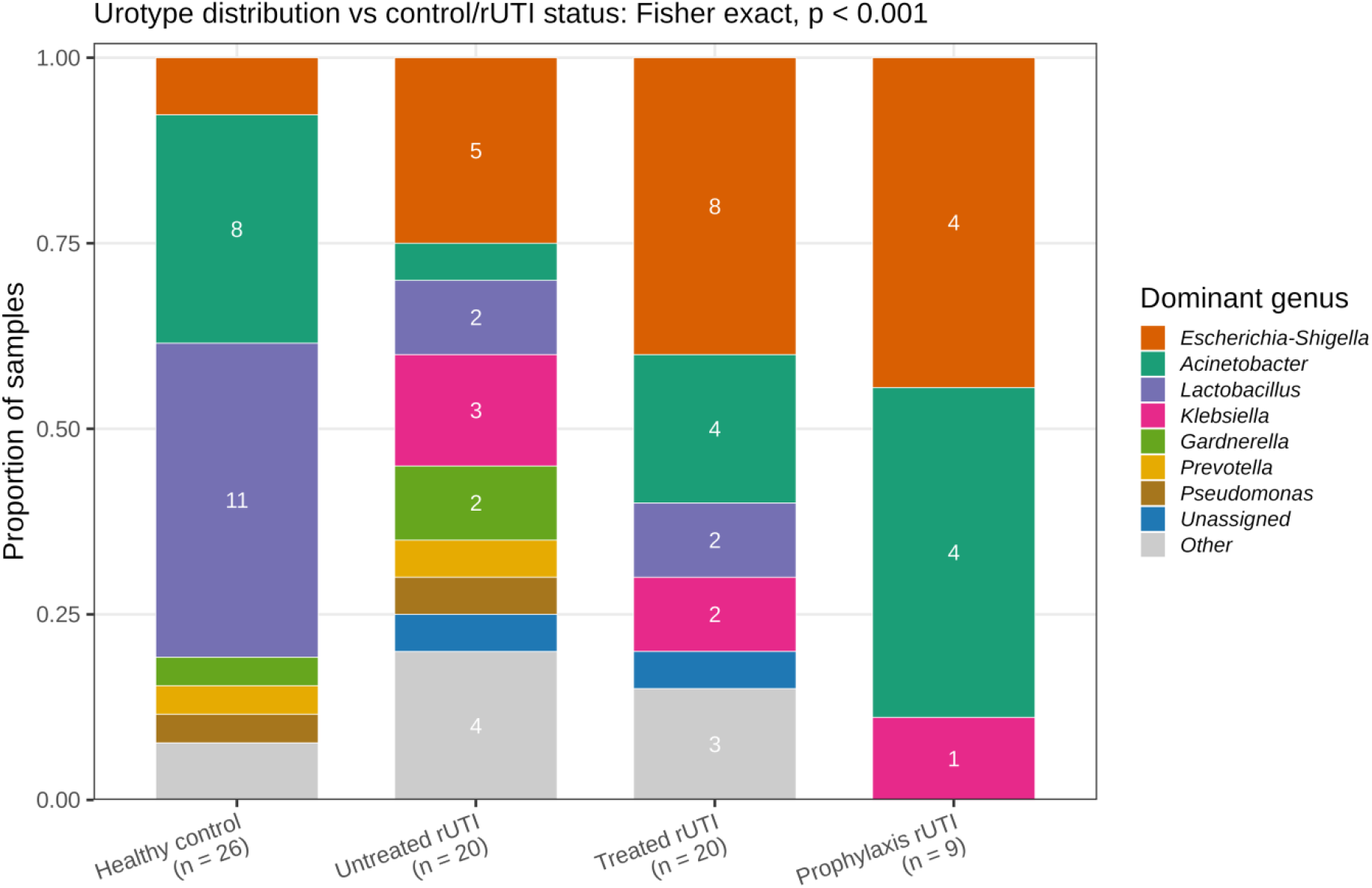
Urotype distribution across study groups. Urotypes were assigned by each sample’s dominant genus; the distribution differed between controls and pooled rUTI (Fisher exact *p* < 0.001). *Lactobacillus*-dominant urotypes predominated in controls (11/26, 42%) but were rare in rUTI (4/49, 8%); *Escherichia*-*Shigella*-dominant urotypes showed the reverse pattern (2/26, 8% vs 17/49, 35%); *Acinetobacter*-dominant urotypes appeared only in the treated (4/20) and prophylaxis (4/9) groups.

